# Contribution of SARS-CoV-2 accessory proteins to viral pathogenicity in K18 hACE2 transgenic mice

**DOI:** 10.1101/2021.03.09.434696

**Authors:** Jesus Silvas, Desarey Morales-Vasquez, Jun-Gyu Park, Kevin Chiem, Jordi B. Torrelles, Roy Neal Platt, Tim Anderson, Chengjin Ye, Luis Martinez-Sobrido

## Abstract

Severe Acute Respiratory Syndrome coronavirus 2 (SARS-CoV-2) is the viral pathogen responsible for the current coronavirus disease 2019 (COVID-19) pandemic. To date, it is estimated that over 113 million individuals have been infected with SARS-CoV-2 and over 2.5 million human deaths have been recorded worldwide. Currently, three vaccines have been approved by the Food and Drug Administration for emergency use only. However much of the pathogenesis observed during SARS-CoV-2 infection remains elusive. To gain insight into the contribution of individual accessory open reading frame (ORF) proteins in SARS-CoV-2 pathogenesis, we used our recently described reverse genetics system approach to successfully engineer recombinant (r)SARS-CoV-2, where we individually removed viral 3a, 6, 7a, 7b, and 8 ORF proteins, and characterized these recombinant viruses *in vitro* and *in vivo*. Our results indicate differences in plaque morphology, with ORF deficient (ΔORF) viruses producing smaller plaques than those of the wild-type (rSARS-CoV-2/WT). However, growth kinetics of ΔORF viruses were like those of rSARS-CoV-2/WT. Interestingly, infection of K18 human angiotensin converting enzyme 2 (hACE2) transgenic mice with the ΔORF rSARS-CoV-2 identified ORF3a and ORF6 as the major contributors of viral pathogenesis, while ΔORF7a, ΔORF7b and ΔORF8 rSARS-CoV-2 induced comparable pathology to rSARS-CoV-2/WT. This study demonstrates the robustness of our reverse genetics system to generate rSARS-CoV-2 and the major role for ORF3a and ORF6 in viral pathogenesis, providing important information for the generation of attenuated forms of SARS-CoV-2 for their implementation as live-attenuated vaccines for the treatment of SARS-CoV-2 infection and associated COVID-19.

**IMPORTANCE:** Despite great efforts put forward worldwide to combat the current coronavirus disease 2019 (COVID-19) pandemic, Severe Acute Respiratory Syndrome coronavirus 2 (SARS-CoV-2) continues to be a human health and socioeconomic threat. Insights into the pathogenesis of SARS-CoV-2 and contribution of viral proteins to disease outcome remains elusive. Our study aims to determine the contribution of SARS-CoV-2 accessory open reading frame (ORF) proteins in viral pathogenesis and disease outcome, and develop a synergistic platform combining our robust reverse genetics system to generate recombinant (r)SARS-CoV-2 with a validated rodent model of infection and disease. We demonstrated that SARS-CoV-2 ORF3a and ORF6 contribute to lung pathology and ultimately disease outcome in K18 hACE2 transgenic mice, while ORF7a, ORF7b, and ORF8 have little impact on disease outcome. Moreover, our combinatory platform serves as the foundation to generate attenuated forms of the virus to develop live-attenuated vaccines for the treatment of SARS-CoV-2.

## INTRODUCTION

Coronaviruses are enveloped, positive sense, single-stranded RNA viruses that cause both mild and lethal upper respiratory illness in humans (1-3). Highly pathogenic and lethal Severe Acute Respiratory Syndrome (SARS-CoV) and Middle Eastern Respiratory Syndrome (MERS-CoV) coronaviruses were first identified in early 2000 and 2010 as spill-over events to humans from horse-shoe bats and dromedary camels, respectively (2, 4, 5). In December 2019, a respiratory infectious disease disseminated throughout the Chinese city of Wuhan and by Spring of 2020, it had spread worldwide (6). A novel coronavirus, Severe Acute Respiratory Syndrome coronavirus-2 (SARS-CoV-2) was identified as the etiological agent of this new coronavirus disease 2019 (COVID-19) pandemic (6-8).

The SARS-CoV-2 genome encodes for 16 non-structural proteins (NSPs) and 6 accessory proteins each encoded by independent open reading frames (ORFs) (9). It has been described that both coronavirus NSP and ORF proteins play important roles in viral replication and transcription, evasion of host immune responses, and viral dissemination (10, 11). In particular, SARS-CoV ORF3a has been implicated as an inducer of membrane rearrangement and cell death (12-14). SARS-CoV ORF3b, ORF6 and nucleocapsid (N) proteins have been described to counteract the host immune interferon (IFN) responses (15). Moreover, SARS-CoV 7a protein has also been shown to inhibit cellular protein synthesis and activation of p38 mitogen-activated protein kinase (16). Conversely, coronavirus ORF7b has been reported to have a Golgi localization signal, where it becomes incorporated into the virion, however no additional studies have delved into any further functions (17, 18). Furthermore, in regard to SARS-CoV-2, early clinical isolates identified a 382-nucleotide deletion leading to a truncated ORF7b and removal of the ORF8 transcription signal (19). Lastly, SARS-CoV-2 ORF8 has been postulated to play a minor role in disease outcome, as natural SARS-CoV-2 isolates containing deletions in the ORF8 has been isolated from individuals presenting COVID-19 (20). Nevertheless, only a limited number of studies have examined the contribution of coronavirus NSP or ORF proteins to pathogenesis and disease outcome in animal models of infection.

To examine the contribution of accessory proteins in the pathogenicity of SARS-CoV-2, we used our previously described reverse genetics approach based on the use of a bacterial artificial chromosome (BAC) to rescue rSARS-CoV-2 with deletions of individual accessory ORF proteins (21). We were able to successfully rescue recombinant (r)SARS-CoV-2 deficient in ORF3a, ORF6, ORF7a, ORF7b, and ORF8 individually. The *in vitro* characterization of each deficient ORF (ΔORF) virus revealed a distinct difference in plaque phenotype in comparison to the recombinant wild-type SARS-CoV-2 (rSARS-CoV-2/WT). This phenotypic change, however, did not affect the viral growth kinetics since all ΔORF rSARS-CoV-2 replicated similarly to rSARS-CoV-2/WT. Interestingly, we observed changes in pathogenesis between the WT and ΔORF rSARS-CoV-2 in the established K18 human angiotensin converting enzyme 2 (hACE2) transgenic mouse model of SARS-CoV-2 infection (22). In contrast to the WT, ORFΔ7a, ORFΔ7b, and ORFΔ8 rSARS-CoV-2, the ORFΔ3a and the ORFΔ6 rSARS-CoV-2 induced less pathology and had a 75% and 50% survival rate, respectively. Furthermore, both ORFΔ3a and ORFΔ6 rSARS-CoV-2 had lower viral titers (10^2^ PFU/ml) at 2 days post-infection (d p.i.), and by 4 d p.i. were no longer detected in nasal turbinates. In contrast, ORFΔ6 viral replication in the lungs reached 10^5^ PFU/ml at 2 d p.i. and only decreased by ∼2-log_10_ at 4 d p.i. ORFΔ3a replication only reached 10^2^ PFU/ml at 2 d p.i. and was not detected by 4 d p.i. in the lungs. Both the ORFΔ7a and ORFΔ7b rSARS-CoV-2 induced similar pathology to rSARS-CoV-2/WT and had a 25% survival rate. By merging our *in vitro* and *in vivo* data, we have been able to generate insights into the contribution of SARS-CoV-2 accessory ORF proteins in the pathogenesis and disease outcome of SARS-CoV-2 infection. These essential data also paved the way for further designing and developing live-attenuated vaccines against SARS-CoV-2.

## RESULTS

### Generation of BACs with deletions in individual accessory ORF proteins

The SARS-CoV-2 genome, which was divided into 5 fragments and chemically synthesized, was assembled into a single BAC that led to efficient virus rescue after transfection into Vero E6 cells using LPF2000 (21). Fragment 1 included the SARS-CoV-2 ORF accessory proteins. Using standard gene engineering approaches, we systematically deleted, individually, ORF3a, ORF6, ORF7a, ORF7b, or ORF8 from fragment 1 using PCR and primer pairs containing BsaI type IIS restriction endonuclease sites. After being confirmed by Sanger sequencing (data not shown), fragment 1 containing the individual deletions in the ORF3a, ORF6, ORF7a, ORF7b, or ORF8 accessory proteins were reassembled into the BAC **(Figure 1)**.

**Figure 1.**
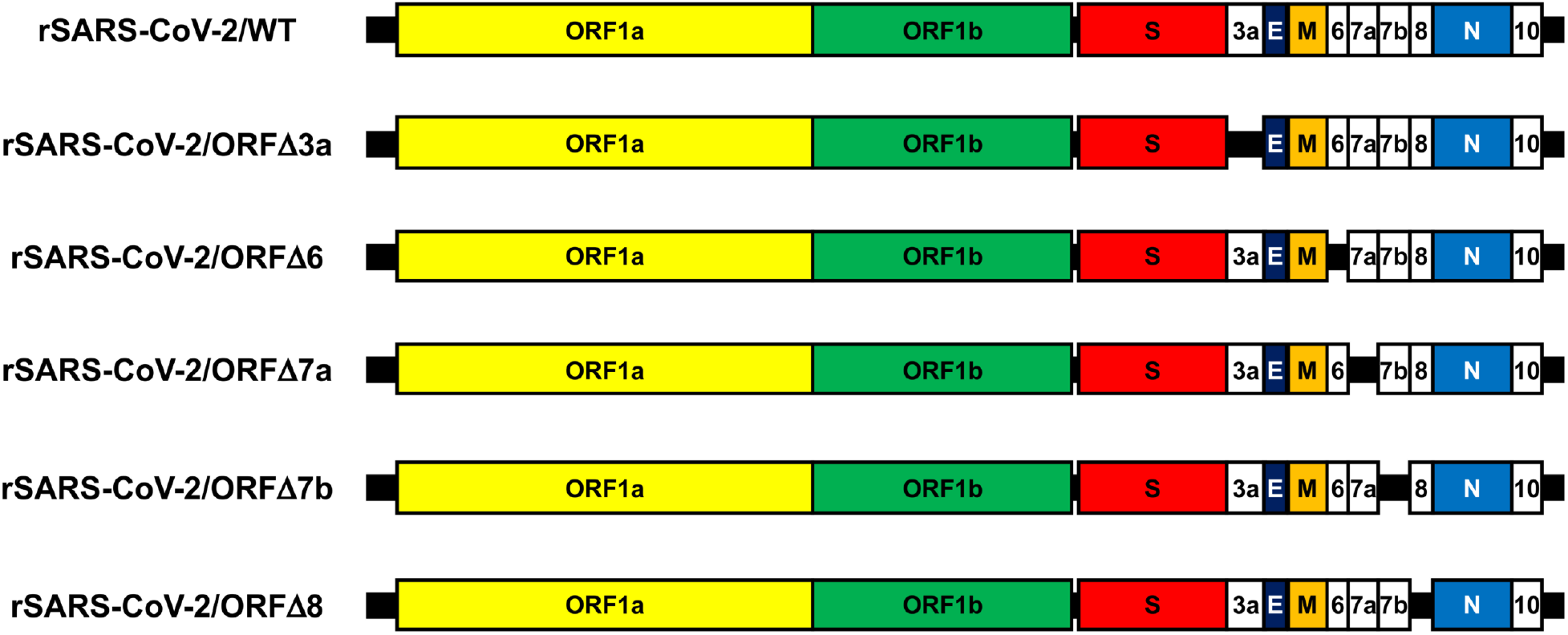
Genome organization of WT and ΔORF rSARS-CoV-2: SARS-CoV-2 genome includes ∼29.8 kb nucleotides among which ∼21.5 kb encodes the ORF1a and ORF1b replicase. The rest of the ∼8.3 kb viral genome encodes the structural spike (S), envelope (E), matrix (M), and nucleocapsid (N) proteins; and the accessory 3a, 6, 7a, 7b, 8, and 10 ORF proteins. Individual deletions of the ORF accessory proteins were introduced into the BAC for rescue of rSARS-CoV-2. Schematic representation not drawn to scale.

### Rescue of ΔORF rSARS-CoV-2

Each BAC with individual deletions in the accessory ORFs were transfected into Vero E6 cells for the recovery of ΔORF rSARS-CoV-2, according to our previously described protocol (21). At 72 h post-transfection, tissue culture supernatants (P0) were collected to inoculate fresh Vero E6 cells (P1). Supernatants were then collected from P1 at 72 hours post-infection (h p.i.) and viral titers, defined as plaque forming units/milliliter (PFU/ml), were determined as previously described (21). To verify rescue of each ΔORF rSARS-CoV-2, indirect immunofluorescence was performed using antibodies directed at the nucleocapsid (N) and spike (S) proteins **(Figure 2A)**. We next verified the individual deletion of each ORF in the rSARS-CoV-2 using RT-PCR procedures to amplify the viral N gene (control), and the regions which cover the corresponding individual ORF deletions **(Figure 2B)**. All the ΔORF rSARS-CoV-2, and rSARS-CoV-2/WT, produced a RT-PCR product of approximately 1.2 kb corresponding to the N gene, whereas amplified regions that cover the corresponding ORF deletions were smaller in the ΔORF rSARS-CoV-2 as compared to rSARS-CoV-2/WT **(Figure 2B)**, demonstrating the deletion of the individual ORFs in the viral genomes. Individual deletions of the viral proteins in the ΔORF rSARS-CoV-2 were further confirmed by Sanger sequencing of PCR products **(Figure 2C)** and by full sequencing of the viral genome (accession number PRJNA707072). These data demonstrate that each ΔORF rSARS-CoV-2 contained the individual deletion of their respective ORF accessory proteins.

**Figure 2.**
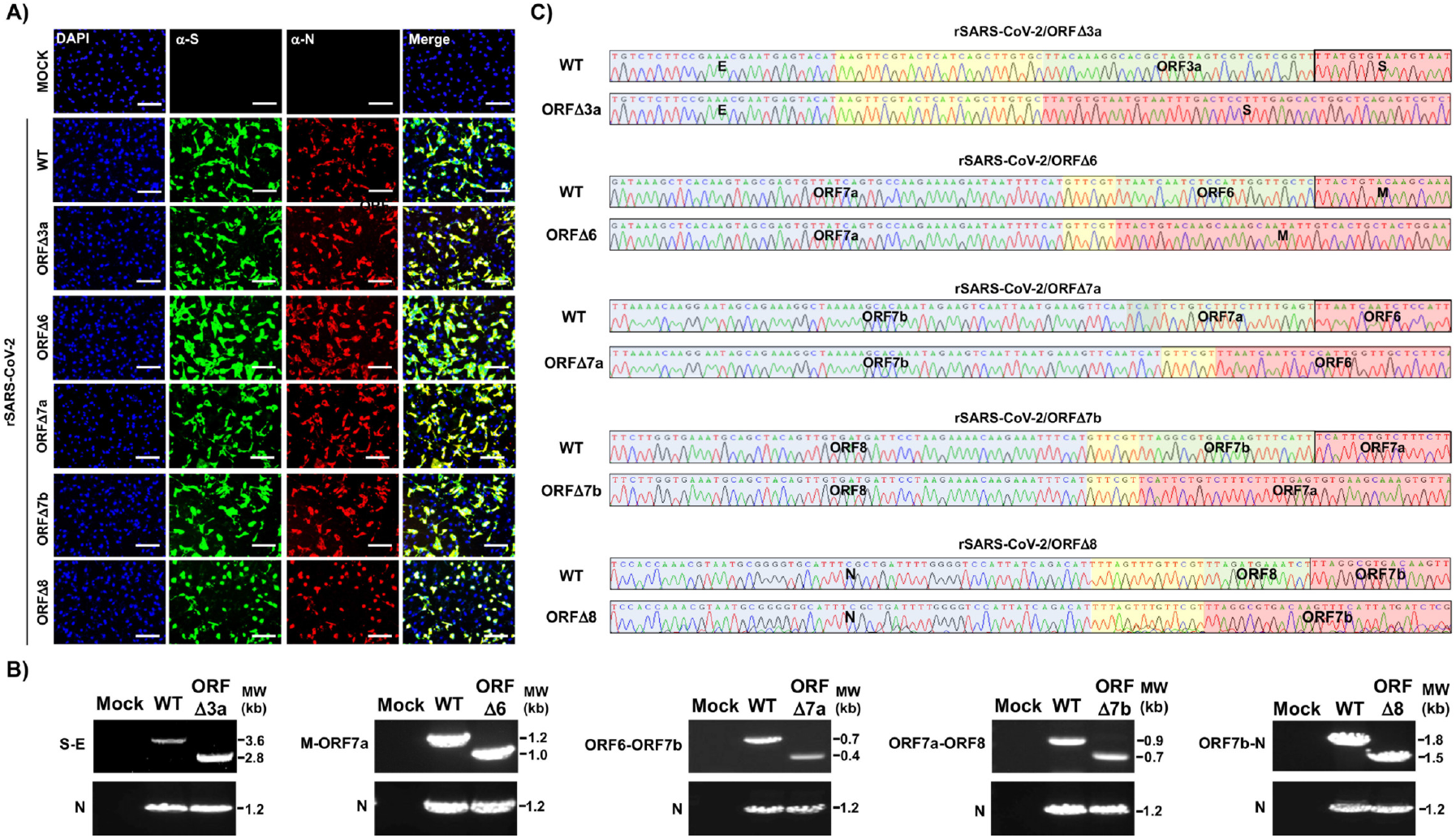
Rescue of ΔORF rSARS-CoV-2. **A**) **IFA:** Vero E6 cells (24-well plate format, 10^5^ cells/well, triplicates) were mock-infected or infected (MOI of 3) with WT, ORFΔ3a, ORFΔ6, ORFΔ7a, ORFΔ7b, or ORFΔ8 rSARS-CoV-2. At 24 h p.i., cells were fixed and immunostained with a cross-reactive polyclonal antibody against SARS-CoV N protein and a cross-reactive monoclonal antibody against SARS-CoV S protein (3B4). Rhodamine red goat anti-rabbit and FITC rabbit anti-mouse secondary antibodies were used and nuclei were visualized by DAPI. Scale bars, 100 μm. **B**) **RT-PCR:** Vero E6 cells (6-well plate format, 10^6^ cells/well) were mock-infected or infected (MOI of 0.1) with WT and ΔORF rSARS-CoV-2 and total RNA was extracted at 24 h p.i. Regions in the viral genome corresponding with the deletion were amplified and the N gene was amplified as an internal control. All the products were separated on a 0.8% agarose gel. **C**) **Sequencing:** RT-PCR products from panel B were gel-purified and subjected to Sanger sequencing. The consensus sequences in the genome of both WT and ΔORF rSARS-CoV-2 downstream the deleted gene are indicated in blue, the intergenic region between viral genes are shown in yellow, the genes deleted in the ΔORF rSARS-CoV-2 are indicated in green, and the viral genes upstream the deleted ORFs in the rSARS-CoV-2 are shown in red.

### Characterization of ΔORF rSARS-CoV-2 *in vitro*

We next proceeded to characterize each ΔORF rSARS-CoV-2 *in vitro* (**Figure 3**). Previous studies have shown that manipulation of the viral genome can affect the phenotype of viral plaques, therefore as an initial step we proceeded to investigate if deletion of individual accessory ORFs had an impact on rSARS-CoV-2 plaque phenotype (23), as some studies have linked changes in plaque phenotype to attenuation or disease outcome (24, 25). Interestingly, we noticed an effect in the plaque phenotype of all ΔORF rSARS-CoV-2 as compared to rSARS-CoV-2/WT in all times p.i. studied (24, 48, 72, and 96 h) **(Figure 3A)**. Next, we compared the growth kinetics of the ΔORF rSARS-CoV-2 to those of the rSARS-CoV-2/WT in Vero E6 cells. To this end, viral titers in the tissue culture supernatants from Vero E6 cells infected (MOI, 0.01) with WT or ΔORF rSARS-CoV-2 collected at 12, 24, 48, 72, and 96 h p.i. were determined by plaque assay **(Figure 3B)**. No statistically significant differences between WT and ORFΔ3a, ORFΔ6, and ORFΔ8 rSARS-CoV-2 were observed at any times p.i., except for the replication of ORFΔ7a and ORFΔ7b rSARS-CoV-2 that was significantly different (∼ 1 log_10_) than those of rSARS-CoV-2/WT at 12 and 24 h p.i. (**Figure 3B**). Peak viral titers for all ΔORF rSARS-CoV-2 and rSARS-CoV-2/WT were observed between 24 and 48 h p.i., with viral titers decreasing (∼ 1 log_10_) at later times p.i., consistent with previous studies with SARS-CoV-2 natural isolates (26, 27). Altogether, these results suggest that despite slight differences in plaque phenotype, ΔORF rSARS-CoV-2 have similar replication kinetics than rSARS-CoV-2/WT in Vero E6 cells.

**Figure 3.**
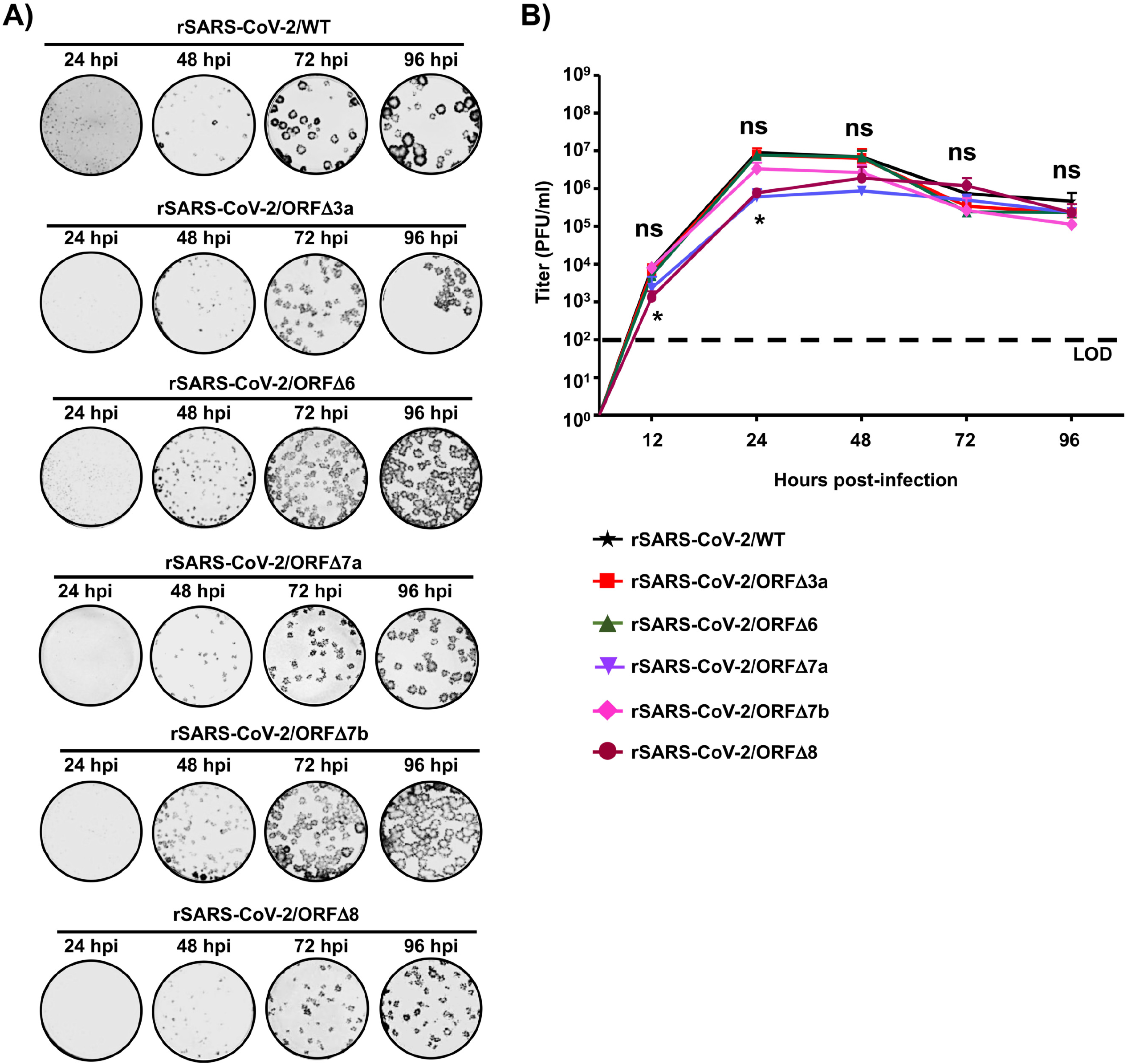
*In vitro* characterization of WT and ΔORF rSARS-CoV-2. **A) Plaque phenotype:** Vero E6 cells (6-well plate format, 10^6^ cells/well) were infected with WT, ORFΔ3a, ORFΔ6, ORFΔ7a, ORFΔ7b, or ORFΔ8 rSARS-CoV-2 and overlaid with media containing agar. Plates were incubated at 37°C and monolayers were immunostained with an anti-N protein SARS-CoV cross-reactive monoclonal antibody 1C7C7 at indicated h p.i. **B) Multicycle growth kinetics:** Vero E6 cells (6-well plate format, 10^6^ cells/well, triplicates) were infected (MOI 0.01) with WT and ΔORF rSARS-CoV-2 and incubated 37°C. At the indicated h p.i. Tissue culture supernatants from infected cells were collected and viral titers were determined by plaque assay (PFU/ml) and immunostaining using the anti-N SARS-CoV cross-reactive monoclonal antibody 1C7C7. Data represent the means ± standard deviations (SDs) of the results determined in triplicate wells. Dotted black lines indicate the limit of detection (LOD, 100 PFU/ml). *P < 0.05: using the Student T test.

### Characterization of ΔORF rSARS-CoV-2 *in vivo*

Coronavirus ORF accessory proteins have been implicated as virulence factors and contribute to both pathogenicity and disease outcome (12, 13, 20, 28-30). Therefore, we proceeded to further investigate the contribution of SARS-CoV-2 ORF3a, ORF6, ORF7a, ORF7b, and ORF8 to viral pathogenicity and disease outcome in our previously established K18 hACE2 transgenic mouse model of SARS-CoV-2 infection and COVID-19 disease (22). Four-to-six-week-old female mice (n=4) were mock (PBS)-infected, infected (10^5^ PFU) with rSARS-CoV-2/WT, or with each of the ΔORF rSARS-CoV-2, and observed for 14 days for morbidity (body weight loss) and mortality (survival) **(Figure 4)**. Our results indicate a similar decrease in body weight percentage up to 5 d p.i., from which ORFΔ3a and ORFΔ7b rSARS-CoV-2 infected mice began to recover **(Figure 4A)**. Interestingly, mice infected with ORFΔ6 and ORFΔ7a rSARS-CoV-2 continued to loss body weight until 7 and 8 d p.i., respectively, and start to recover **(Figure 4A)**. All mice infected with WT or ORFΔ8 rSARS-CoV-2 succumbed to viral infection by 6 and 7 d p.i., respectively **(Figure 4B)**. Our continued observations of mice infected with ORFΔ3a, ORFΔ6a, ORFΔ7a, and ORFΔ7b rSARS-CoV-2 identified survival rates of 75%, 50%, 25%, and 25%, respectively **(Figure 4B)**. Upon full comparison, it is worth noting that despite the increased morbidity observed in mice infected with the ORFΔ3a rSARS-CoV-2, 3 out of 4 mice survived viral infection, suggesting ORF3a could play an important role in viral pathogenesis.

**Figure 4.**
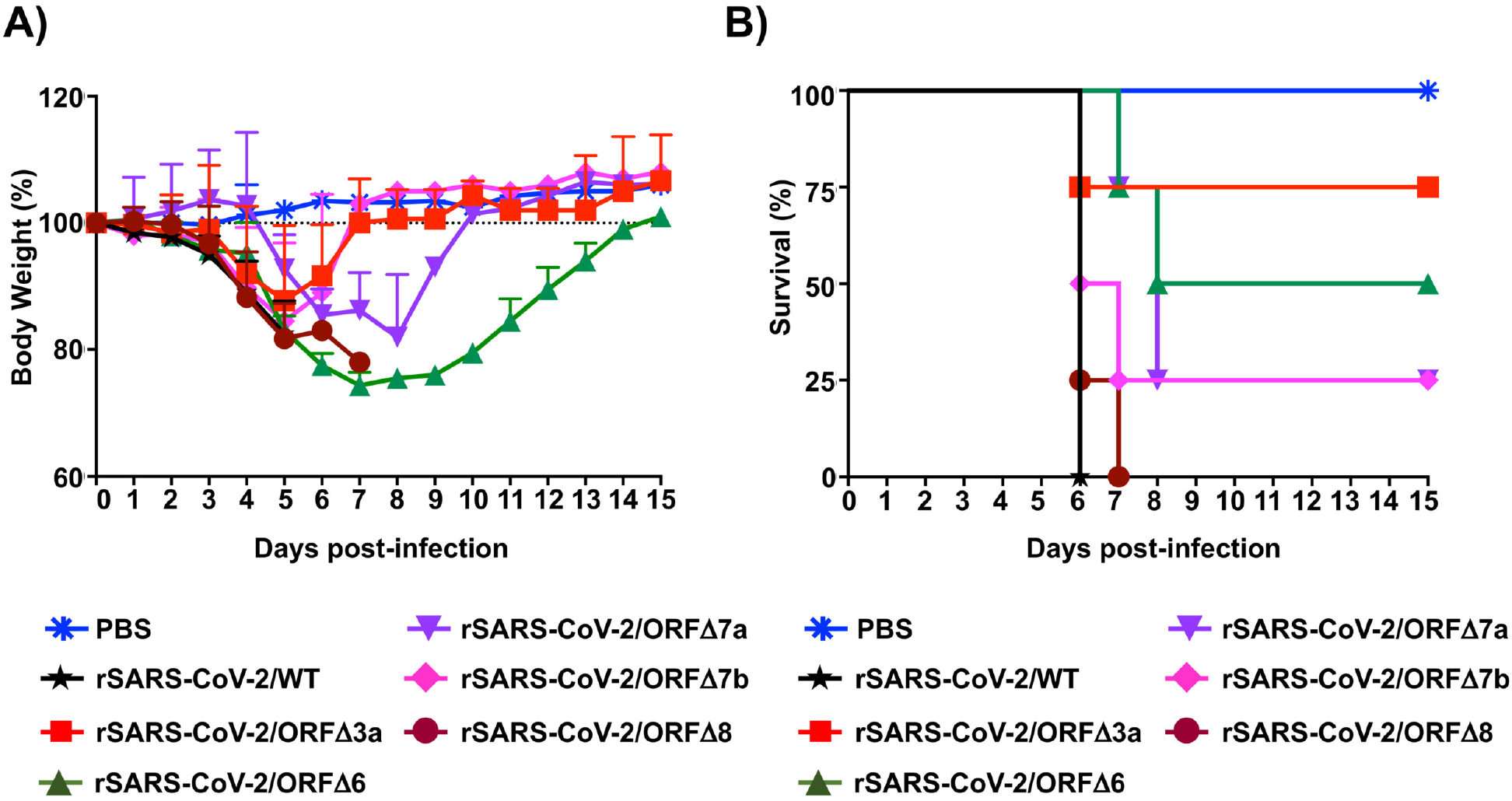
Infection of K18 hACE2 transgenic mice with WT and ΔORF rSARS-CoV-2: K18 hACE2 transgenic female 6-8-week-old mice were mock (PBS)-infected or infected (i.n.) with 10^5^ PFU of WT and ΔORF rSARS-CoV-2 (n=4/group). (**A)** Body weight and (**B)** survival were evaluated at the indicated d p.i. Mice that lost >25% of their initial body weight were humanely euthanized. Error bars represent SDs of the mean for each group.

We next evaluated viral replication in nasal turbinates and lungs of K18 hACE2 transgenic mice infected with the WT or ΔORF rSARS-CoV-2 at 2 and 4 d p.i. **(Figure 5)**. At 2 d p.i. WT and ORFΔ6 rSARS-CoV-2 were detected in nasal turbinates in all mice, with rSARS-CoV-2/WT reaching titers of up to 5×10^3^ PFU/ml, while ORFΔ6 rSARS-CoV-2 peaked at 5×10^2^ PFU/ml. We only detected ORFΔ3a rSARS-CoV-2 (10^2^ PFU/ml) in 50% of infected mice while ORFΔ7b rSARS-CoV-2 replicated up to 3×10^3^ PFU/ml in 75% of infected mice. Only 50% of mice had detectable levels (range between 0.5-1×10^3^ PFU/ml) of ORFΔ8 rSARS-CoV-2 **(Figure 5A)**. Interestingly, out of all the ΔORF rSARS-CoV-2 tested, ORFΔ7a replicated in the nasal turbinate to levels (5×10^4^ PFU/ml) higher than those observed with rSARS-CoV-2/WT. By 4 d p.i. ORFΔ3a and ORFΔ6 rSARS-CoV-2 were no longer detected in nasal turbinates, while 75% of mice infected with rSARS-CoV-2/WT had viral titers ranging between 5×10^1^ to 5×10^2^ PFU/ml. ORFΔ7a, ORFΔ7b and ORFΔ8 rSARS-CoV-2 replicated to lower levels (∼5×10^1^ PFU/ml) than rSARS-CoV-2/WT **(Figure 5A)**. In the lungs WT, ORFΔ6 and ORFΔ7a rSARS-CoV-2 were detected at levels of ∼10^5^ PFU/ml at 2 d p.i. (**Figure 5B**). Viral titers of ORFΔ7b and ORFΔ8 (∼10^4^ PFU/ml) and ORFΔ3a (∼0.5×10^2^ PFU/ml) rSARS-CoV-2 were lower than those of rSARS-CoV-2/WT (**Figure 5B**). Interestingly, by 4 d p.i., ORFΔ3a rSARS-CoV-2 was no longer detected in the lungs while WT, ORFΔ6, ORFΔ7a, ORFΔ7b, and ORFΔ8 rSARS-CoV-2 only decreased ∼1 log_10_ to those observed by 2 d p.i. **(Figure 5B)**.

**Figure 5.**
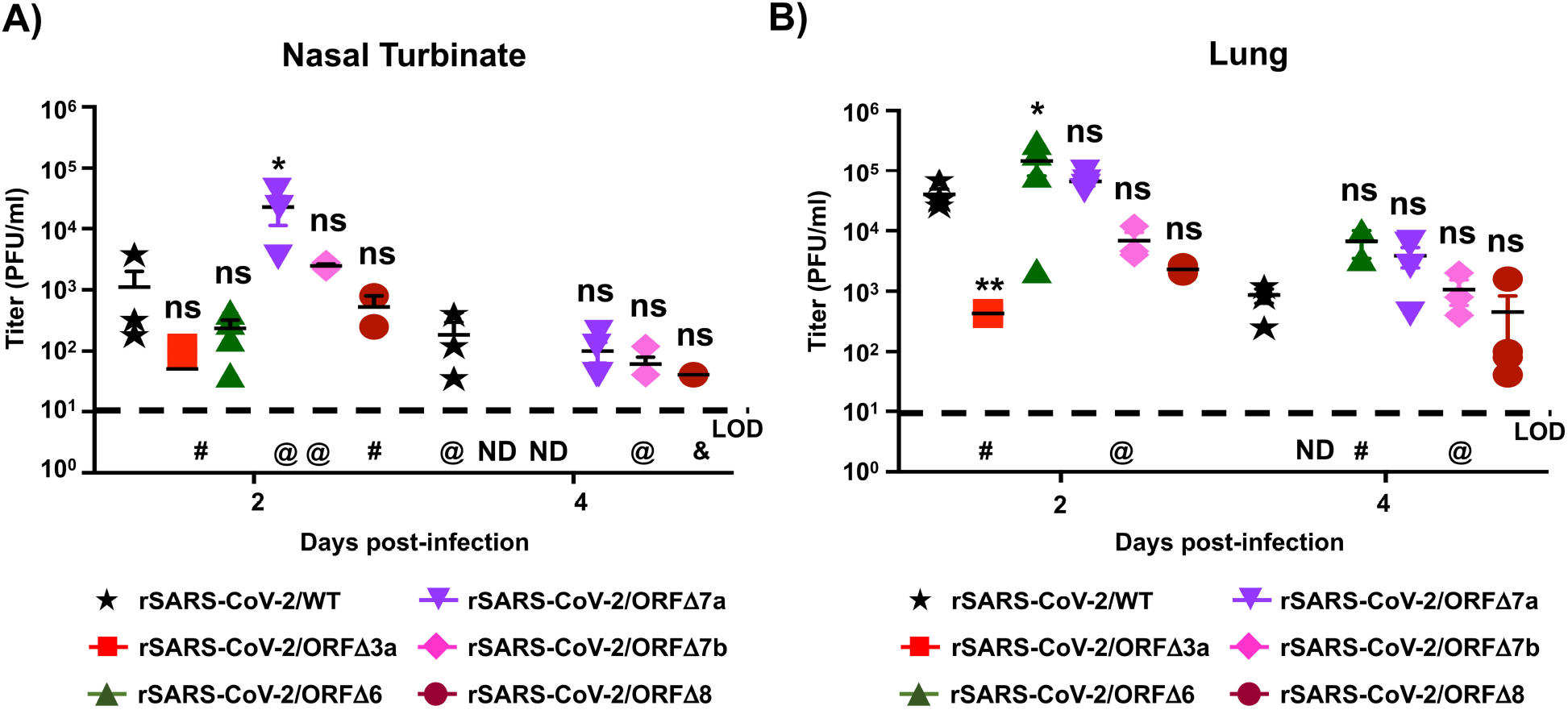
Replication of WT and ΔORF rSARS-CoV-2 in nasal turbinate and lungs: K18 hACE2 transgenic female 6-8-week-old mice were mock (PBS)-infected or infected (i.n.) with 10^5^ PFU of WT and ΔORF rSARS-CoV-2 (n=8/group). Mice were sacrificed at 2 (n=4/group) and 4 (n=4/group) d p.i. and viral titers nasal turbinate and lung were determined by plaque assay (PFU/ml) and immunostaining using the cross-reactive SARS-CoV 1C7C7 N protein monoclonal antibody. Viral titers in the nasal turbinate **(A)** and lungs **(B)** are shown. Symbols represent data from individual mouse, and bars the geometric means of viral titers. Dotted lines indicate limit of detection (LOD, 10 PFU/ml). ND, not detected; @, not detected in 1 mouse; #, not detected in 2 mice: &, not detected in 3 mice. Negative results of the PBS-infected mice are not plotted.

Lastly, to evaluate the impact of viral infection in the lungs of infected animals, we performed gross pathology analysis on lungs collected at 2 and 4 d p.i. (**Figure 6A**). Both WT and ΔORF rSARS-CoV-2 induced similar pathology at 2 d p.i., with WT, ORFΔ7a, and ORFΔ7b rSARS-CoV-2 inducing pathological lesions in more than 50% of lung area **(Figure 6B)**. Intriguingly, by 4 d p.i. only WT and ORFΔ8 rSARS-CoV-2-infected lungs maintained pathological lesions, while ORFΔ3a, ORFΔ6, ORFΔ7a, ORFΔ7b rSARS-CoV-2 infected lungs appear to recover from lesions observed at 2 d p.i. Altogether, these *in vivo* results provide a new insight into the contribution of SARS-CoV-2 accessory ORF proteins in viral pathogenesis and disease, suggesting a major role of ORF3a and ORF6 and a less impact of ORF7a, ORF7b, and ORF8 in virulence and disease outcome.

**Figure 6.**
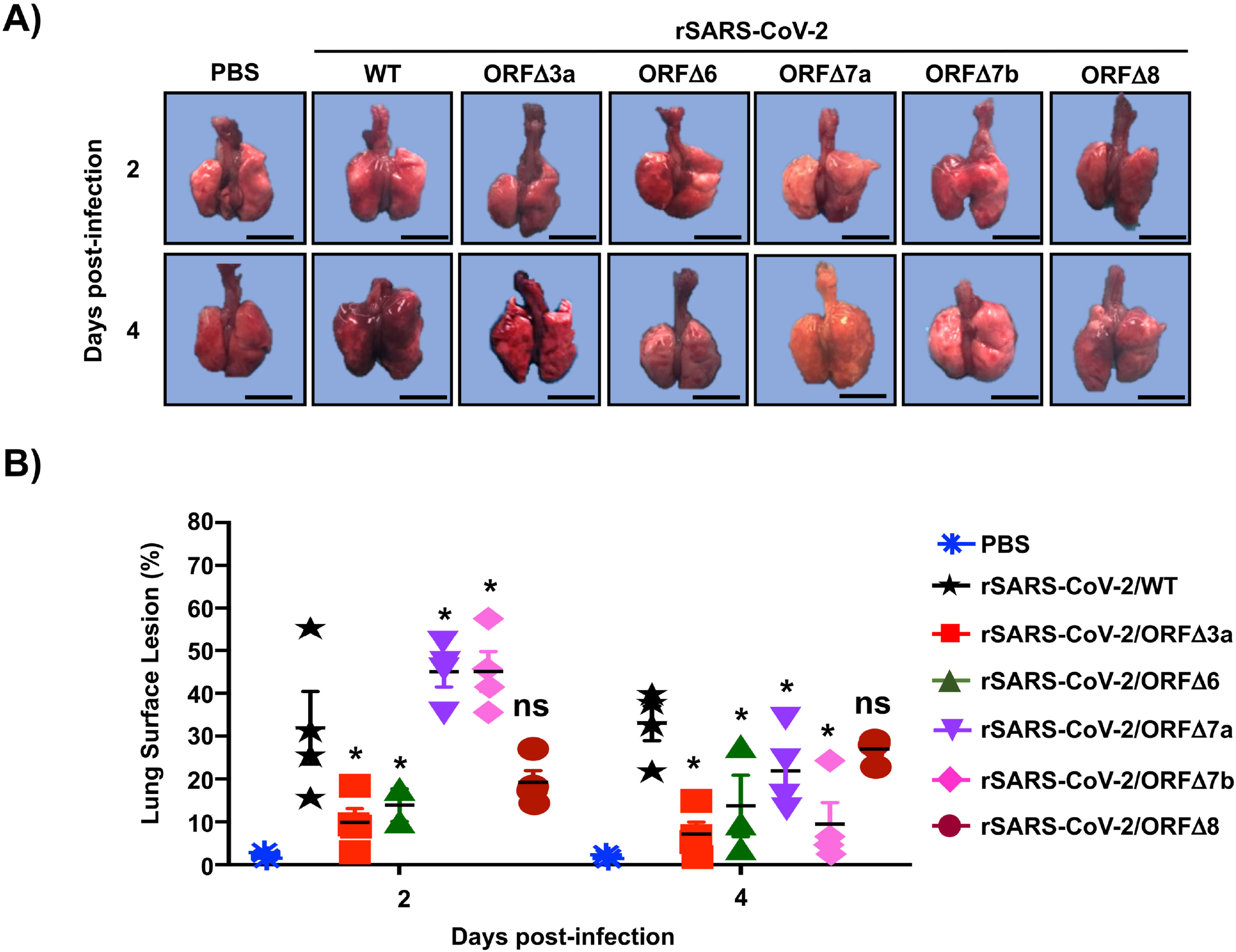
Gross pathology analysis of lungs from K18 hACE2 transgenic mice infected with WT and ΔORF rSARS-CoV-2: Lungs from 6-to-8-week-old female K18 hACE2 transgenic mice (n=8/group) mock (PBS)-infected or infected (i.n.) with 10^5^ PFU of WT or ΔORF rSARS-CoV-2 were harvested at 2 (n=4/group) or 4 (n=4/group) d p.i. and imaged **(A)**, and scored for viral induced lesions **(B)**. Calculation of total lung surface area affected by viral induced lesions. Scale bars, 3 cm.

## DISCUSSION

Viral proteins contribute to different aspects of disease outcome and pathogenesis during the course of viral infections (20, 28, 29). Reverse genetics systems allow for the deletion of viral proteins to gain insight into their contribution to viral replication, pathogenesis, transmission, and/or in disease outcome, among others (31-34). We have recently established a *state-of-the-art* reverse genetics system for the efficient rescue of rSARS-CoV-2/WT (21, 22) that we have used to generate ΔORF rSARS-CoV-2 and interrogate the contribution of the ORF3a, ORF6, ORF7a, ORF7b, and ORF8 accessory proteins in viral replication *in vitro* and *in vivo*. Similar studies have provided insight into the contribution of accessory ORF proteins in SARS-CoV infection *in vitro*, and have identified a variety of functions, ranging from inhibition of host immune responses to activation of host cell death pathways (12-14, 17, 30, 31, 35-37). However, to date, no study has provided insight into the role of SARS-CoV-2 ORF accessory proteins in viral replication *in vitro* and/or in pathogenesis *in vivo*. Our study using robust genetics systems and the validated K18 hACE2 transgenic mouse model of SARS-CoV-2 infection provides us with useful information on the contribution of SARS-CoV-2 ORFs 3a, 6, 7a, 7b, and 8 accessory proteins to viral pathogenesis.

In this study, we generated rSARS-CoV-2 deficient in ORFs 3a, 6, 7a, 7b, or 8 accessory proteins **(Figure 1)** and characterized them both *in vitro* **(Figures 2 and 3)** and *in vivo* **(Figures 4-6)**. Importantly, we have been able to demonstrate the ORF deficient nature of these rSARS-CoV-2 (**Figure 2**). During our initial *in vitro* characterization, we first identified that ORF proteins contributed to early dissemination and formation of detectable viral plaques in Vero E6 cell monolayers: viruses lacking 3a, 7a, 7b, and 8 ORF proteins developed smaller plaques than rSARS-CoV-2/WT (**Figure 3**). This was a surprising finding, as we only observed ∼1 log_10_ difference in growth kinetics between the WT and any of the ΔORF rSARS-CoV-2 (**Figure 3**).

To further determine the contribution of each ORF accessory protein in SARS-CoV-2 pathogenesis, we used our recently established K18 hACE2 transgenic mouse model (**Figure 4**) (22). This particular *in vivo* study is the first, to our knowledge, that analyzes the contribution of SARS-CoV-2 ORF accessory proteins in viral infection in a validated animal model of infection. We observed a broad range of morbidity and survival outcomes between each of the ΔORF rSARS-CoV-2 as compared to rSARS-CoV-2/WT (**Figure 4**). SARS-CoV-2 ORF accessory proteins are encoded in the following order: 3a, 6, 7a, 7b, and 8. Our data collectively showed greatest survival in mice infected with ORFΔ3a (75%) and 100% mortality with the ORFΔ8 rSARS-CoV-2. In this regard, a natural SARS-CoV-2 variant with a deletion in the ORF8 has been recently isolated from patients presenting COVID-19 symptoms (20). Thus, our results with the ORFΔ8 rSARS-CoV-2 in the K18 hACE2 transgenic mouse model correspond to those observed in people infected with a natural ORFΔ8 SARS-CoV-2 isolate (20). Our findings with ORFΔ3a and ORFΔ6 rSARS-CoV-2 warrant further characterization as there is limited insight into mechanistic exploitation of host pathways by ORF3a (12, 13, 30, 35, 36). Since it is well known that SARS-CoV-2 ORF6 is a potent inhibitor of the host innate immune response (11, 29), we were expecting low viral load and an increased immune response; however, since SARS-CoV N protein also inhibits host immune responses (15, 38), it is plausible that SARS-CoV-2 N protein may have a similar function and is responsible of counteracting host innate immune and inflammatory responses. Even though our studies did not analyze cytokine and chemokine production, the gradual recovery of mice infected with ORFΔ6 rSARS-CoV-2 are suggestive of viral clearance over time. Analysis of innate and adaptive immune responses would be important, as well as modulation of IFN secretion, ion signaling channels, and cellular apoptotic and/or necrosis pathways, to determine the contribution of these ORF accessory proteins in viral pathogenesis *in vivo*. This is currently the focus of our ongoing *in vivo* studies with these ΔORF rSARS-CoV-2. Overall, similar to what was described with rSARS-CoV, our study highlights that SARS-CoV ORFs 3a, 3b, 6, 7a, and 7b had no significant impact on viral replication *in vivo* (31), but ORF 3a seems to be involved in virulence, as its absence decreases SARS-CoV-2 virulence.

Overall, this study demonstrates the robustness of our BAC-based reverse genetics approach to generate rSARS-CoV-2, including those with deletions of ORF accessory proteins, and provides information on the contribution of ORFs 3a, 6, 7a, 7b, and 8 accessory proteins in viral fitness *in vitro* (Vero E6 cells) and *in vivo*, in our recently established K18 hACE2 transgenic mouse model of SARS-CoV-2 infection and COVID-19 disease (22). Importantly, information from this study also provides novel insights for the generation of attenuated forms of SARS-CoV-2 for the development of live-attenuated vaccines for the treatment of this important respiratory pathogen, and its associated COVID-19 disease.

## MATERIALS AND METHODS

### Biosafety

All the *in vitro* and *in vivo* experiments with infectious natural isolate or rSARS-CoV-2 were conducted under appropriated biosafety level (BSL) 3 and animal BSL3 (ABSL3) laboratories, respectively, at Texas Biomedical Research Institute. Experiments were approved by the Texas Biomed Institutional Biosafety (IBC) and Animal Care and Use (IACUC) committees.

### Cells

African green monkey kidney epithelial cells (Vero E6, CRL-1586) were obtained from the American Type Culture Collection (ATCC, Bethesda, MD) and maintained in Dulbecco’s modified Eagle medium (DMEM) supplemented with 5% (v/v) fetal bovine serum, FBS (VWR) and 100 units/ml penicillin-streptomycin (Corning).

### Reverse genetics system

The BAC harboring the entire viral genome of SARS-CoV-2 USA-WA1/2020 (Accession No. MN985325) was described previously (21). Deletion of individual accessory ORF proteins was achieved in viral fragment 1 by using inverse PCR and primer pairs containing a BsaI type IIS restriction endonuclease site. All the primer sequences are available upon request. Fragments including the individual deletion of accessory ORF proteins were reassembled into the BAC using BamHI and RsrII restriction endonucleases.

### Rescue of ΔORF rSARS-CoV-2

Virus rescues were performed as previously described (21). Briefly, confluent monolayers of Vero E6 cells (10^6^ cells/well, 6-well plates, triplicates) were transfected with 4.0 μg/well of SARS-CoV-2 BAC using Lipofectamine 2000. After 24 h, transfection media was exchanged for p.i. media [DMEM supplemented with 2% (v/v) FBS], and cells were split and seeded into T75 flasks 48 h post-transfection. After incubation for another 72 h, tissue culture supernatants were collected, labeled as P0 and stored at -80°C. After viral titration of the supernatant, the P0 virus was used to infect fresh Vero E6 cells at multiplicity of infection (MOI) of 0.0001 to make new viral stocks. Tissue culture supernatants were collected 72 h p.i., aliquoted, labeled as P1, and stored at -80°C for future use.

### Immunofluorescence assay (IFA)

Vero E6 cells (10^5^ cells/well, 24-well plate format, triplicates) were mock-infected or infected (MOI of 3) with WT or ΔORF rSARS-CoV-2. At 24 h p.i, cells were fixed with 10% formaldehyde solution at 4°C overnight and permeabilized using 0.5% (v/v) Triton X-100 in PBS for 15 min at room temperature. Cells were incubated overnight with 1 μg/ml of a SARS-CoV N protein cross-reactive polyclonal antibody at 4°C and a SARS-CoV-2 spike (S) cross-reactive monoclonal antibody (3B4), washed with PBS, and stained with a FITC-labeled goat anti-mouse IgG (1:200) and Rhodamine-labeled goat anti-rabbit IgG (1:200). Nuclei were visualized by DAPI staining. After washing with PBS, cells were visualized and imaged with an EVOS microscope (ThermoFisher Scientific).

### Plaque assay and immunostaining

Confluent monolayers of Vero E6 cells (10^6^ cells/well, 6-well plate format, triplicates) were infected with serial diluted viruses for 1 h at 37°C. After viral adsorption, cells were overlaid with p.i. media containing 1% low melting agar and incubated at 37°C. At 72 h p.i., cells were fixed overnight with 10% formaldehyde solution. For immunostaining, cells were permeabilized with 0.5% (v/v) Triton X-100 in PBS for 15 min at room temperature and immunostained using the SARS-CoV N protein cross-reactive monoclonal antibody 1C7C7 (1 μg/ml) and the Vectastain ABC kit (Vector Laboratories), following the manufacturers’ instruction. After immunostaining, plates were visualized on a Chemi-Doc (Bio-Rad).

### Virus growth kinetics

Confluent monolayers of Vero E6 cells (10^6^ cells/well, 6-well plate format, triplicates) were infected (MOI of 0.01) with WT or ΔORF rSARS-CoV-2. After 1 h virus adsorption at 37°C, cells were washed with PBS and incubated in p.i. media at 37°C. At the indicated times after infection, viral titers in tissue culture supernatants were determined by plaque assay and immunostaining using the SARS-CoV N protein cross-reactive monoclonal antibody 1C7C7, as previously described (39).

### Reverse transcription-polymerase chain reaction (RT-PCR)

Total RNA from Vero E6 cells (10^6^ cells/well, 6-well plate format) infected (MOI of 0.01) with WT or ΔORF rSARS-CoV-2 was extracted with TRIzol Reagent (Thermo Fisher Scientific) according to the manufacturer’s instructions. RT-PCR amplification was performed using Super Script II Reverse transcriptase (Thermo Fisher Scientific) and Expanded High Fidelity PCR System (Sigma Aldrich). The amplified DNA products were subjected to 0.7% agarose gel analysis and the gel-purified PCR fragments were subjected to sanger sequencing (ACGT). All primer sequences used for RT-PCR are available upon request. Methods for Illumina library preparation, sequencing, and analysis of recombinant viruses follow those previously described (21).

### Mice

Specific-pathogen-free, 4–8 week-old, female B6.Cg-Tg(K18-ACE2)2Prlmn/J (Stock No: 034860) K18 hACE2 transgenic mice were purchased from The Jackson Laboratory (Bar Harbor, ME). Our previous studies did not show significant sex dependent differences in morbidity, mortality and viral titers (22). K18 hACE2 transgenic mice were maintained in micro-isolator cages at ABSL-3 and provided sterile water and chow *ad libitum* and acclimatized for one week prior to experimental manipulation. For morbidity and mortality studies, a total of 20 females were used (n=4/group), whereas a total of 40 female mice (n=4 per group) were used for viral titers at 2 and 4 d p.i.. For the mock infected control groups in the morbidity and mortality studies, 2 female K18 hACE2 transgenic mice were used and a total of 4 (n=2 per time-point), were used as mock-infected controls for gross pathology analysis.

### Mouse infection and sample processing

K18 hACE2 transgenic mice were either mock (PBS)-infected or infected intranasally (i.n.) with 10^5^ PFU of WT or ΔORF rSARS-CoV-2 in a final volume of 50 µL following isoflurane sedation. After viral infection, mice were monitored daily for 14 days for morbidity (body weight loss) and mortality (survival). Mice showing >25% loss of their initial body weight were defined as reaching experimental endpoint and humanely euthanized. For viral titers and gross pathology analysis, K18 hACE2 transgenic mice were infected as above but euthanized at 2 or 4 d p.i. Nasal turbinates and lung tissues were harvested and homogenized in 1 ml of PBS using a Precellys tissue homogenizer (Bertin Instruments) for viral titrations. Tissue homogenates were centrifuged at 21,500 × *g* for 5 min and supernatants were collected for determination of viral titer.

### Statistical analysis

Statistical analysis was performed using GraphPad Prism 8.3. For multiple comparisons, a 2-way ANOVA with multiple comparisons was performed.

## ACKNOWLEDGEMENTS

We want to thank Dr. Thomas Moran at the Icahn School of Medicine at Mount Sinai for providing us with the SARS-CoV cross-reactive N monoclonal antibody 1C7C7. We also want to thank BEI Resources for providing the SARS-CoV-2 USA-WA1/2020 isolate (NR-52281). Finally, we would also like to thank members at our institutes for their efforts in keeping them fully operational during the COVID-19 pandemic and the BSC and IACUC committees for reviewing our protocols in a time efficient manner. We would like to dedicate this manuscript to all COVID-19 victims and to all heroes battling this disease.

